# Heritable tumor cell division rate heterogeneity induces clonal dominance

**DOI:** 10.1101/097683

**Authors:** Margriet M. Palm, Marjet Elemans, Joost B. Beltman

## Abstract

Tumors consist of a hierarchical population of cells that differ in their phenotype and genotype. This hierarchical organization of cells means that a few clones (i.e., cells and several generations of offspring) are abundant while most are rare, which is called *clonal dominance*. Such dominance also occurred in published *in vitro* iterated growth and passage experiments with tumor cells in which genetic barcodes were used for lineage tracing. A potential source for such heterogeneity is that dominant clones derive from cancer stem cells with an unlimited self-renewal capacity. Furthermore, ongoing evolution within the growing population may also induce clonal dominance. To understand how clonal dominance developed in the iterated growth and passage experiments, we built a computational model that accurately simulates these experiments. The model simulations reproduced the clonal dominance that developed in *in vitro* iterated growth and passage experiments when the division rates vary between cells, due to a combination of initial variation and of ongoing mutational processes. In contrast, the experimental results can neither be reproduced with a model that considers random growth and passage, nor with a model based on cancer stem cells. Altogether, our model suggests that *in vitro* clonal dominance develops due to selection of fast-dividing clones.

## Introduction

Intratumoral heterogeneity, the genotypic and phenotypic differences within a single tumor, is a well known feature of cancer [1] and strongly influences the effectiveness of cancer therapy [2]. Genotypic heterogeneity is the result of random mutations, and while most of these mutations are neutral passenger mutations, some are functional mutations that add to phenotypic heterogeneity. Phenotypic differences may also be caused by phenomena such as differential signaling from the local tumor micro-environment, epigenetic changes, and stochastic gene expression [3]. Another proposed source of intratumoral, phenotypic heterogeneity is the presence of so-called *cancer stem cells* (CSCs) that have an unlimited potential to renew and can give rise to *di erentiated cells* (DCs) with a limited potential to renew [4]. The presence of CSCs would result in a tumor containing a mixture of CSCs, and DCs that all derive from a small number of CSCs.

For a long time, evidence for the presence of CSCs was primarily based on xenograft models in which transplantation of tumor cells into immunodeficient mice resulted in tumor growth in only a small fraction of the mice [1, 5], suggesting that only a subset of the tumor cells has the ability to sustain long-term growth. However, the lack of success of initiating tumor growth in immunodeficient mice may also be related to the incomplete inhibition of the immune response [6], or to the dramatic change in tumor micro-environment upon transplantation [5]. An alternative approach to identify the existence of CSCs is to perform *lineage tracing* by fluorescent marking of a subpopulation of cells [7, 8]. For example, Schepers *et al.* [9] managed to trace the lineage of CSCs by fluorescently labeling cells expressing stem cell markers in mice developing intestinal adenomas and thereby showed that all cells in small adenomas descended from a single stem cell. However, fluorescent labeling of stem cells is not possible in all cancer types, and for those cancer types an alternative approach is taken by labeling a small fraction of the tumor cells in animal models. Studies employing this strategy showed that the number of colored patches reduced during tumor growth, while the size of these patches increased [10–12]. These observations are compatible with the hypothesis that tumor cells descend from a small number of CSCs.

Another, high-resolution, approach to lineage tracing is the application of unique genetic tags, also called *cellular barcodes*, to a population of tumor cells [13–19]. Tumors grown in immunodeficient mice injected with barcoded tumor cells are dominated by cells that express only a small subset of the barcodes [15, 16]. Serial implantation of barcoded leukemic cells showed that rare clones in one mouse can develop dominance after transplantation into a second mouse, indicating that clonal dominance is not a predetermined property of certain clones [18]. Porter *et al.* [13] used cellular barcoding to follow the development of clonal dominance over time in an *in vitro* setup, thereby controlling the external factors that could affect clonal dominance. Populations derived from several polyclonal cell lines were barcoded and grown for three days after which a fraction of the cells was passed on to the next generation. By repeating this process (Fig 1A) and analyzing the intermediate clone distributions, Porter *et al.* [13] showed that clonal dominance progressed over time (Fig 1B). Interestingly, similar experiments with a monoclonal K562 cell line resulted in a minimal progression of clonal dominance, hinting that an intrinsic variability in the cell population may cause the progression of clonal dominance. However, Nolan-Stevaux *et al.* [15] did not observe a strong development of clonal dominance during 8 days of *in vitro* growth without passaging, indicating that random passage could also play a role in developing clonal dominance.

Although the studies discussed above represent a strong base of evidence for the development of clonal dominance *in vivo* as well as in *in vitro* tumor cell populations, the mechanism that drives this dominance remains unknown. The presence of CSCs is consistent with the induction of clonal dominance, but only in some cancers direct evidence for the involvement of CSCs is available [5]. Alternatively, evolution may cause clones with a higher division or survival rate to dominate the tumor. Hence, it is necessary to further investigate the role of these mechanism in the development of clonal dominance. One way to do this in a formal way is to construct computational models that incorporate different hypotheses and compare the outcome of computer simulations to spatio-temporal clonal dynamics observed experimentally. Such an approach has been used before in numerous studies addressing the temporal and spatial evolution of tumor cell populations with CSCs, which are thoroughly reviewed in [20]. Several of these modeling studies focused on the development of spatial tumor heterogeneity by employing cell-based models in which intratumoral heterogeneity is induced by CSCs [21, 22], by CSCs and epigenetic changes [23], or by mutations [24]. Here, we built a computational model that simulates the iterated growth and passage experiments described in [13]. By incorporating different hypotheses for the development of clonal dominance in our model, we show that heritable variability of the rates of cell divisions amongst tumor cells, which is due to a combination of initial variation and mutational processes, is su cient to induce clonal dominance in iterated growth and passage experiments.

## Results

### Re-analysis of published sequencing data

**Figure 1:**
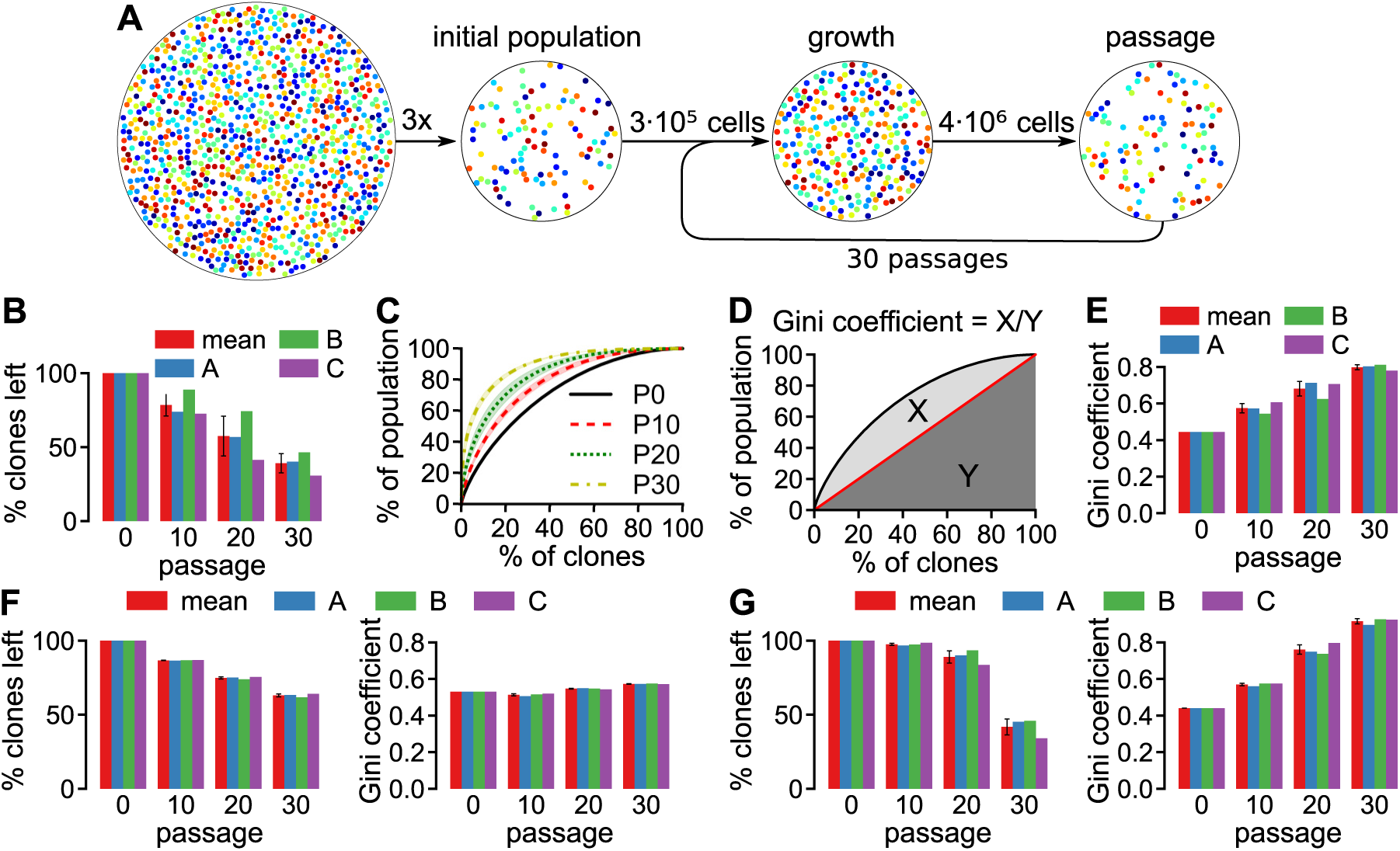
Setup and results of the in vitro iterated growth and passage experiment previously described by Porter et al. [13]. A Experimental setup. B-E Development of the clone size distribution of polyclonal K562 cells, as obtained from our own analysis of the FASTQ files published by Porter *et al.* [13]. Shown are the percentage of clones that remain after each passage (B), the percentage of clones versus the percentage of the population taken up by those clones (C, mean ± SD and 3 biological replicates shown) and the Gini coe cient (E; ratio of areas X and Y in D). F Clone loss (left) and Gini coe cient (right) for the *in vitro* experiments with the monoclonal K562 cell line. G Clone loss (left) and Gini coe cient (right) for the *in vitro* experiments with HeLa cells. All error bars depict the SD of the 3 replicates.

We started by re-analyzing the data for the *in vitro* experiments with the lentivirally barcoded cell lines, in order to subsequently make quantitative comparisons with simulation results (see Methods section for more details). In these previously published experiments, the barcode-transduced cells were grown and aliquots containing 3 • 10^5^ cells were used to initialize three biological replicates of the iterated growth and passage experiment. The cells were then, iteratively, grown for 3 days after which 3 • 10^5^ cells were passed on to the next generation (Fig 1**A**).

In agreement with Porter *et al.* [13], iterated growth and passage with the polyclonal K562 cell line resulted in clone loss (Fig 1**B**) and progressing clonal dominance (Fig 1**C**). To evaluate clonal dominance, we plotted clone size, sorted from large to small, against the cumulative population fraction for clones of that size (Fig 1**C**). In the remainder of this paper, we quantify clonal dominance based on this graph (1D) by employing the *Gini coe cient* [25]:

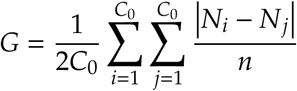

with *C_0_* the number of clones at initialization, *N_i_* the size of clone *i*, and *n* the number of cells. As expected from the observed clone distributions, the Gini coe cient is already greater than zero at the start of the experiment (indicating mild clonal dominance) and further increases over time (Fig 1**E**). We also processed the data for the *in vitro* experiments with monoclonal K562 cells (Fig 1**F**) and with HeLa cells (Fig 1**G**) and clone loss was in good agreement with those published in [13] (S1 Fig). The monoclonal K562 cell line exhibits a more limited clone loss and development of clonal dominance (at time points beyond the initial measurement) compared to the polyclonal K562 cell line. The HeLa cells show a similar trend as the K562 cells: the number of unique clones declines while clonal dominance increases. However, clone loss occurred later for the HeLa cells than it did for the polyclonal K562 cells.

### Iterated growth and passage reduces the number of clones, but does not cause progressive clonal dominance

The limited development of clonal dominance during passaging in monoclonal K562 cells compared to polyclonal K562 cells indicates that chance could play a role in the development of clonal dominance, hence the most straightforward explanation for the development of clonal dominance is that the iterated passages cause small clones to completely disappear while larger clones remain and grow. This hypothesis is supported by Porter’s *in vivo* experiments in which no passage occurred and no loss of clones nor clonal dominance was observed [13]. To test whether clone loss during passage explains clone loss and progressive clonal dominance, Porter *et al.* employed a computational model of iterated deterministic growth phases and passage steps in which a subset of the cells were selected at random, showing that this could partially explain the experimentally observed clone loss but not the development of clonal dominance (see Methods section of [13]). In these simulations, all clones grow according to *N i* (*t* +*t*_growth_) = *N i* (*t*)*e^rt^*^growth^, where *N_i_(t)* represents the size of clone *i* at time *t*, *t*growth is the duration of the growth phase, and *r* is the division rate of the cells. At the end of the passage interval, *n*_pass_ cells are selected randomly and passed to the next generation (Fig 1**A**). As a first step, we confirmed the Porter simulation results using the code that was published with Porter’s paper [13] (https://github.com/adaptivegenome/clusterseq/). We ran simulations starting with *C* =14,000 clones uniformly distributed over 3 • 10^5^ cells and 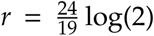 to match the experimentally observed doubling time of 19 hours. As expected, these simulations resulted in a moderate reduction of the number of clones (Fig 2A; blue bars). In contrast to the *in vitro* results, clonal dominance developed only slightly and hardly increased over time beyond passage 10 (Fig 2B; blue bars). To test if a realistic initial clone distribution improved the resemblance between the experimental observations and the simulation results, we extended the simulations by employing the initial clone distribution of the polyclonal K562 cell line (Fig 2C) in which clonal dominance is already slightly developed. In this setting the expected number of clones left still remained larger than that for the polyclonal K562 cells while the clone distribution was hardly a ected (Fig 2A-B; green bars).

**Figure 2:**
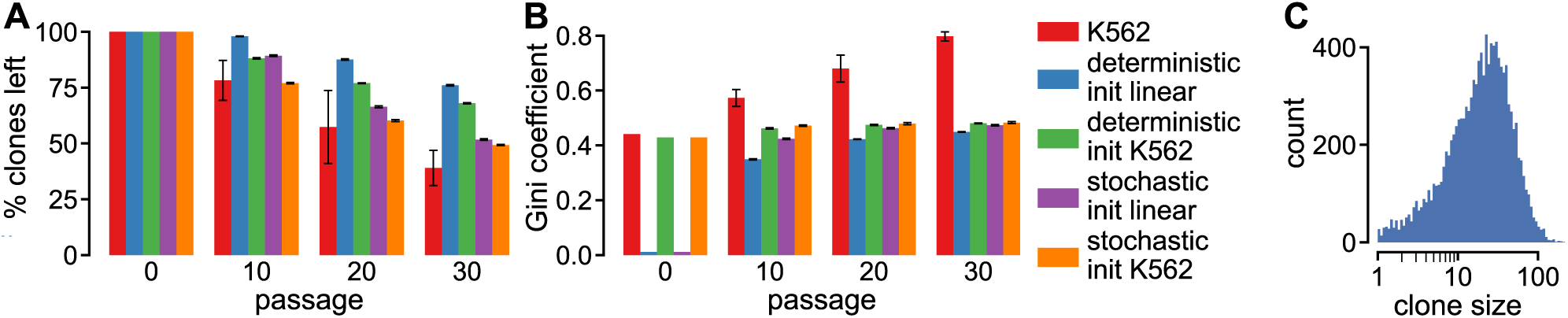
Clonal dominance does not develop during passaging of cells that divide at a fixed rate. A-B Clone loss (A) and Gini coe cient (B) during iterated growth and passage with either deterministic or stochastic growth and initialized either with a uniform clone size distribution or the initial distribution for polyclonal K562 cells. All values are the mean of 10 simulations and the error bars represent the SD. C histogram of the initial clone sizes of polyclonal K562 cells.

Porter *et al.* [13] noted that in simulations in which more cells were passaged, fewer clones disappeared, indicating that the random process of passage causes some clones to become smaller and finally disappear. However, because cell division is modeled as a deterministic process, the clone sizes, relative to other clones, remain similar over time, and the probability to disappear for individual clones remains unchanged over time. The assumption of deterministic growth would be reasonable for large clones, but for small clones, probabilistic events could strongly affect the simulation outcome. As shown in Fig 2C, a large part of the clones contain only a few cells, e.g., Ȉ 25% of the clones have less than 10 cells. Therefore, we next explored the e ect of stochastic cell division by replacing deterministic growth by a stochastic growth model using Gillespie’s Stochastic Simulation Algorithm (SSA) [26] (see Methods section for details and Table 1 for the model parameters). In contrast with Porter’s deterministic growth model, passage occurs when the population size reaches the critical population size *n* crit = 4 • 10^6^, which corresponds to approximately 3 days of growth with a population doubling time of 19 hours. Because growth continues until the critical population size is reached, the division rate no longer a ects the simulation results and we therefore set it to the arbitrary value of 1. As before, *n*_pass_ cells are then chosen randomly and passed on to the next generation.

We performed stochastic growth simulations initialized with either a uniform clone distribution (Fig 2A-B; purple bars) or the initially observed clone distribution of polyclonal K562 cells (orange bars). In both cases the results were similar to those with deterministic growth, albeit with a larger decrease in the number of clones. For simulations with stochastic growth that were initialized with the polyclonal K562 clone distribution, the clone loss resembles the average *in vitro* clone loss for passages 10 and 20, but for passage 30 the *in vitro* clone loss overtook the simulated clone loss. This suggests that the early clone loss is dominated by the effects of stochastic division and passage, and that at a later stage another, unknown, mechanism further increases clone loss. Altogether, these results indicate that passage and growth can cause clone loss during passage, but neither of these mechanisms can induce progressive clonal dominance.

**Table 1:** Parameters of the stochastic growth and passage models.

| parameter | symbol | default value | unit |
| --- | --- | --- | --- |
| *Iterated growth & passage* |  |  |  |
| population size after passage | $n_{\text{pass}}$ | $3 \cdot 10^5$ | cells |
| maximum population size | $n_{\text{crit}}$ | $4 \cdot 10^6$ | cells |
| number of passages |  | 30 |  |
| *Tau leaping algorithm* |  |  |  |
| tau leap step | $\tau$ | .0005 | days |
| *Stochastic growth model* |  |  |  |
| division rate | $r$ | 1 | day<sup>-1</sup> |
| *CSC growth model* |  |  |  |
| symmetric division probability | $p_1$ | 0.5 | |
| asymmetric division probability | $p_2$ | 0.5 | |
| symmetric commitment probability | $p_3$ | 0 | |
| initial CSC fraction | $\text{CSC}_0$ | 5% | |
| maximum DC age | $M$ | 10 | divisions |
| CSC division rate | $r_{\text{CSC}}$ | 1 | day<sup>-1</sup> |
| DC division rate | $r_{\text{DC}}$ | $\frac{24}{19}$ | day<sup>-1</sup> |
| *Agent-based model* |  |  |  |
| mean initial division rate | $r_0$ | 1 | day<sup>-1</sup> |
| initial division rate SD | $\sigma_0^*$ | 0 | |
| mutation SD | $\sigma_m^*$ | 0 | |

### The presence of CSCs does not induce clonal dominance

The presence of cancer stem cells is thought to induce tumor heterogeneity because they generate a, hierarchically organized, population of cancer stem cells and di erentiated cells [27]. To introduce cancer stem cells in our growth model we replaced the unlimited growth in the stochastic model with a previously published model of cancer stem cell driven growth [28]. In this model cells are either cancer stem cells (CSCs) that can divide indefinitely, or di erentiated cells (DCs) that divide a limited number of times. CSCs proliferate at a division rate of *r*CSC and division can result either in two stem cells (with probability *p_1_*), in a stem cell and a di erentiated cell (with probability *p_2_*), or in two di erentiated cells (with probability *p_3_*) (Fig 3A). Di erentiated cells proliferate at a division rate *r*DC until they reach their maximum number of divisions *M*, after which, following Weekes *et al.* [28], they die with a rate *r*DC. We did not consider random cell death of cancer stem cells and di erentiated cells, because this process only a ects the population growth rate.

**Figure 3:**
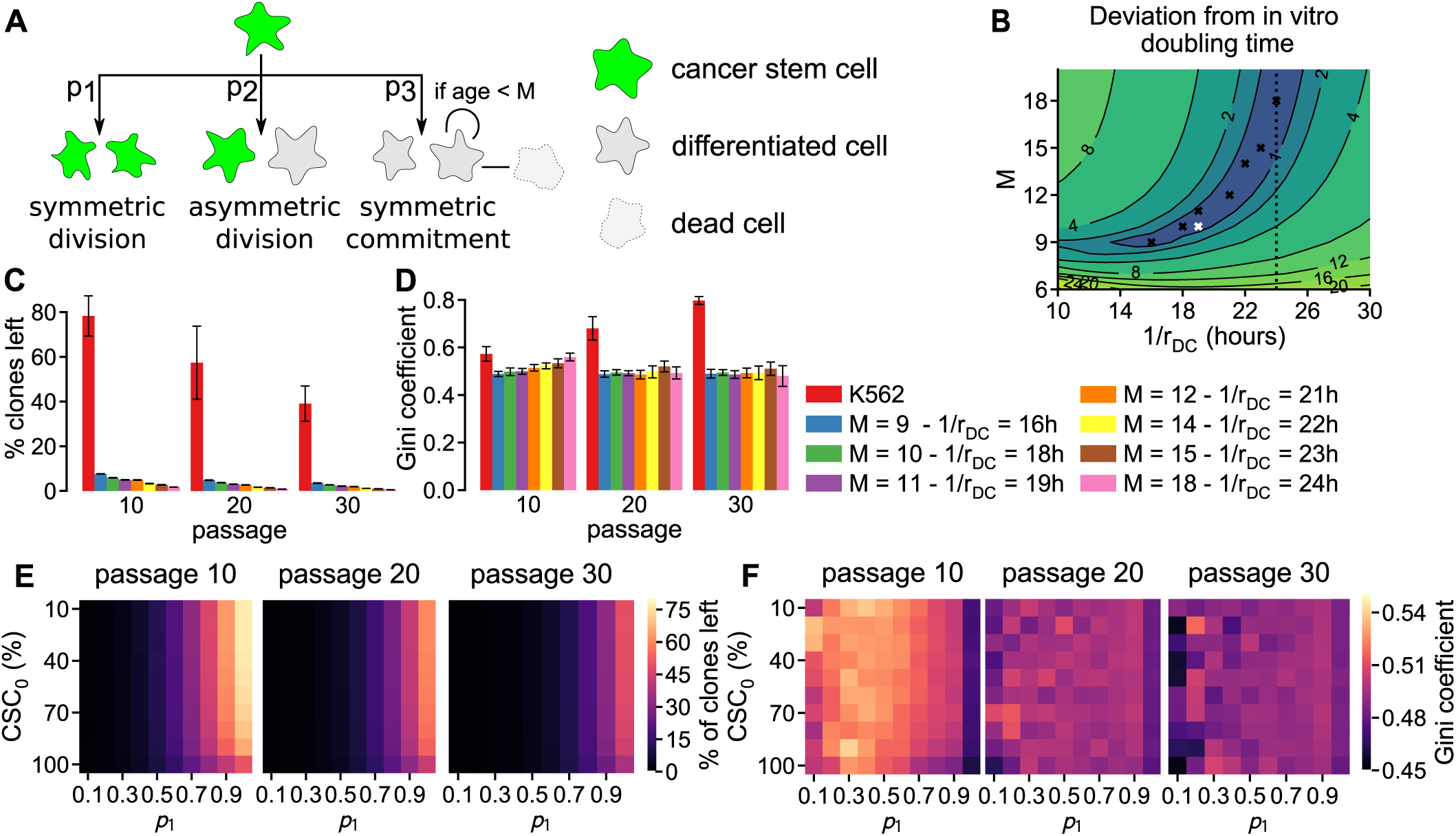
Simulations with CSC model result in massive clone loss and no development of clonal dominance. A Scheme illustrating the divisions and cell death in the CSC growth model. B Heatmap showing the di erence between *in vitro* and simulated population doubling time (19 hours) depending on the maximum number of DC divisions (*M*) and DC division rates (*r*DC) in the CSC growth model. The white cross denotes the default model settings and the black crosses depict several alternative parameter sets that result in a similar population doubling time. C-D Clone loss (C) and Gini coe cient (D) for the parameter sets highlighted in B and all other parameters as in Table 1. E-F E ect of symmetric CSC division probability (*p_1_*) and initial CSC percentage (CSC_0_) on clone loss (E) and Gini coe cient (F), with all other parameters as in Table 1. All values are the mean of 10 simulation replicates with the error bars depicting the SD.

We fine-tuned the parameters of the CSC growth model such that the population growth rate is consistent with the 19 hour doubling time reported by Porter *et al.* [13]. For this we exploited the analytical solution of the CSC model elegantly derived by Weekes *et al.* [28]. This solution predicts that the population of cells initially grows, and then develops according to one of three growth regimes determined by *β* = (*p* 1 – *p* 3)*r*_CSC_. When *β* > 0, the population continues to grow, when *β* = 0 the population reaches an equilibrium, and when < 0 the population will eventually go extinct. Because the *in vitro* cells were reported to be in “*log phase growth*” [13], we limited the parameter space to *β* > 0 by setting *p_3_* to 0, which ensures a monotonically growing cell population. A complicating factor is that the parameters determining do not a ect the growth in the first couple of days. Instead, during this time interval, growth is determined by the division rate of DCs (*r*DC) relative to that of CSCs (*r*DC), and the maximum age of DCs (*M*). Therefore, we explored how much the simulated population doubling time deviates from the experimental doubling time of 19 hours when *M* and *r*DC were varied while the other simulation parameters remained at their default value according to Table 1 (Fig 3B). This exploration resulted in a well-defined parameter range for which the *in vitro* doubling time could be reproduced *in silico*.

To illustrate the clonal development of the CSC growth model, we tested the model for several parameter sets from the region that matches with the experimentally observed population growth rate (Fig 3B; crosses). The simulations showed that the maximum age of DCs or the CSC division rate hardly a ect clonal development (Fig 3C-D). Furthermore, an at first sight surprisingly strong reduction in clone number occurred in the simulations while the clone distribution remained virtually unchanged (Fig 3C-D). The dramatic reduction in clone number can readily be explained: clones frequently loose all CSCs during passage, which takes away the long-term self-renewal capacity of such clones. We tested this explanation by varying the initial percentage of CSCs and the probability of a symmetric CSC di erentiation while keeping the other parameters constant (Table 1 and white cross in Fig 3B). In line with our explanation for the massive clone loss, increasing the probability of symmetric CSC division strongly reduced clone loss (Fig 3E). Increasing the initial percentage of stem cells had a less pronounced e ect on clone loss (Fig 3E), which fits with the observation in [28] that the proportion of CSCs in the population will become constant in the long term and does not depend on the initial percentage of CSCs. Altogether, increasing the probability of a symmetric CSC to 1 and the initial CSC percentage to 100% did not lead to clone loss matching that observed experimentally. Moreover, changing these parameters had only little effect on clonal dominance (Fig 3F). The lack of progression of clonal dominance over time can be explained by a lack in difference between the clones: although in the long run all cells descend from only a few clones, there is no difference in the speed at which each clone generates offspring. In conclusion, incorporating CSC growth into our passaging simulation resulted in a poor match to the experimental observations because far too many clones disappeared and the distribution of the remaining clones did not exhibit clonal dominance.

### A model with evolving division rates results in clone loss and increasing clonal dominance

Because neither simulations with random growth and passaging of tumor cells nor simulations incorporating CSC growth can explain the development of clonal dominance, we next considered intrinsic variability of growth characteristics as a source of clonal dominance development. To this purpose, we let go of the distinction between cell types and let all cells divide with their own division rate *r i*, that is subject to mutations. Note that the term mutation is used here to describe any change in a cell’s phenotype that can be inherited, which includes both genetic and epigenetic changes. Mutation of cellular properties requires the explicit representation of single cells, therefore we can no longer use Gillespie’s SSA because clones rather than single cells are explicitly described in that type of simulation. Therefore, we implemented an agent based model (ABM) in which each cell is represented explicitly, and both a barcode and division rate are associated to each cell.

In the ABM there are two sources for division rate variability. First, the division rate of cells at initialization may vary due to prior mutations. To implement this, the division rate of each cell is initialized to *r_i_* = *X_i_r*_0_, with *r_i_* the cell’s division rate, *r*_0_ the mean division rate of all cells and *X_i_* taken from a normal distribution with mean 1 and SD 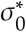. Second, mutations are included by setting the division rate of each o spring to *r_i_* = *Y_i_r_p_*, with *Y_i_* taken from a normal distribution with mean 1 and SD 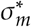, and *r_p_* the division rate of the parent cell. Further details on the implementation of the ABM can be found in the Methods.

**Figure 4:**
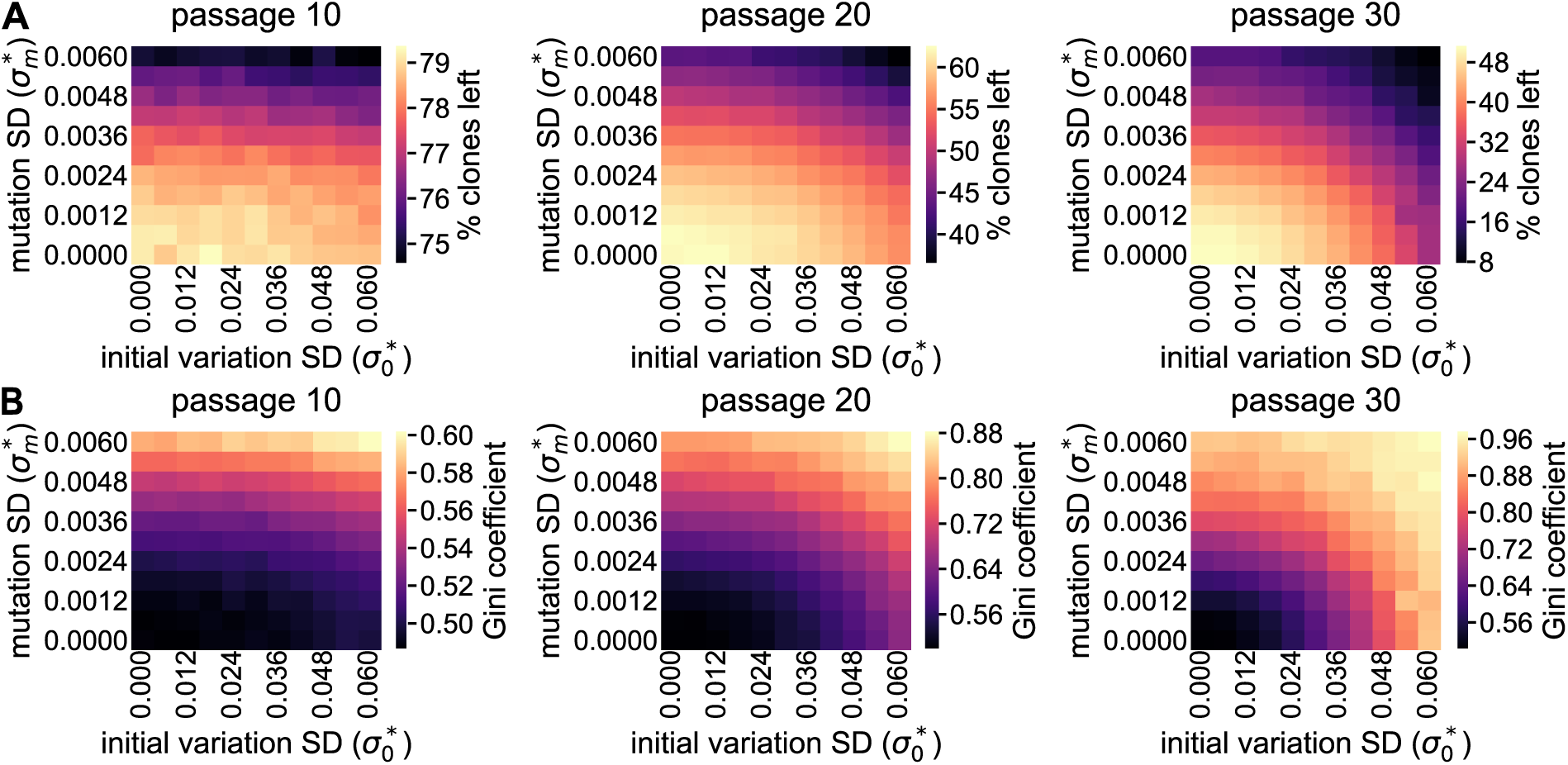
The ABM that describes evolution of division rate variability induces clone loss and clonal dominance A-B. Clone loss (A) and Gini coe cient (B) for a range of initial division rate SDs (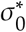) and mutation SDs (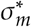), with all other parameters as in Table 1 and all data points representing the mean for 10 simulations.

To test if the ABM can induce similar clone loss (Fig 4A) and clonal dominance (Fig 4B) as observed *in vitro*, we ran simulations for a range of initial division rate SDs (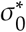) and mutation SDs (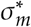), all initialized using the initial distribution of the polyclonal K562 cells. Note that we did not vary the initial mean division rate (*r*_0_) because this parameter does not a ect the simulation results (S2 Fig). The resulting ranges of clone loss and Gini coe cient do indeed include those observed for the polyclonal K562 cells. Furthermore, simulations with division rates that vary due to either initial variation (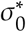 > 0) or mutations (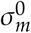) result in higher levels of clone loss and a higher Gini coe cient compared to simulations without any variation (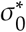 = 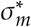). Thus, the ABM that describes evolution of division rate variability has the potential to match the *in vitro* results.

**Figure 5:**
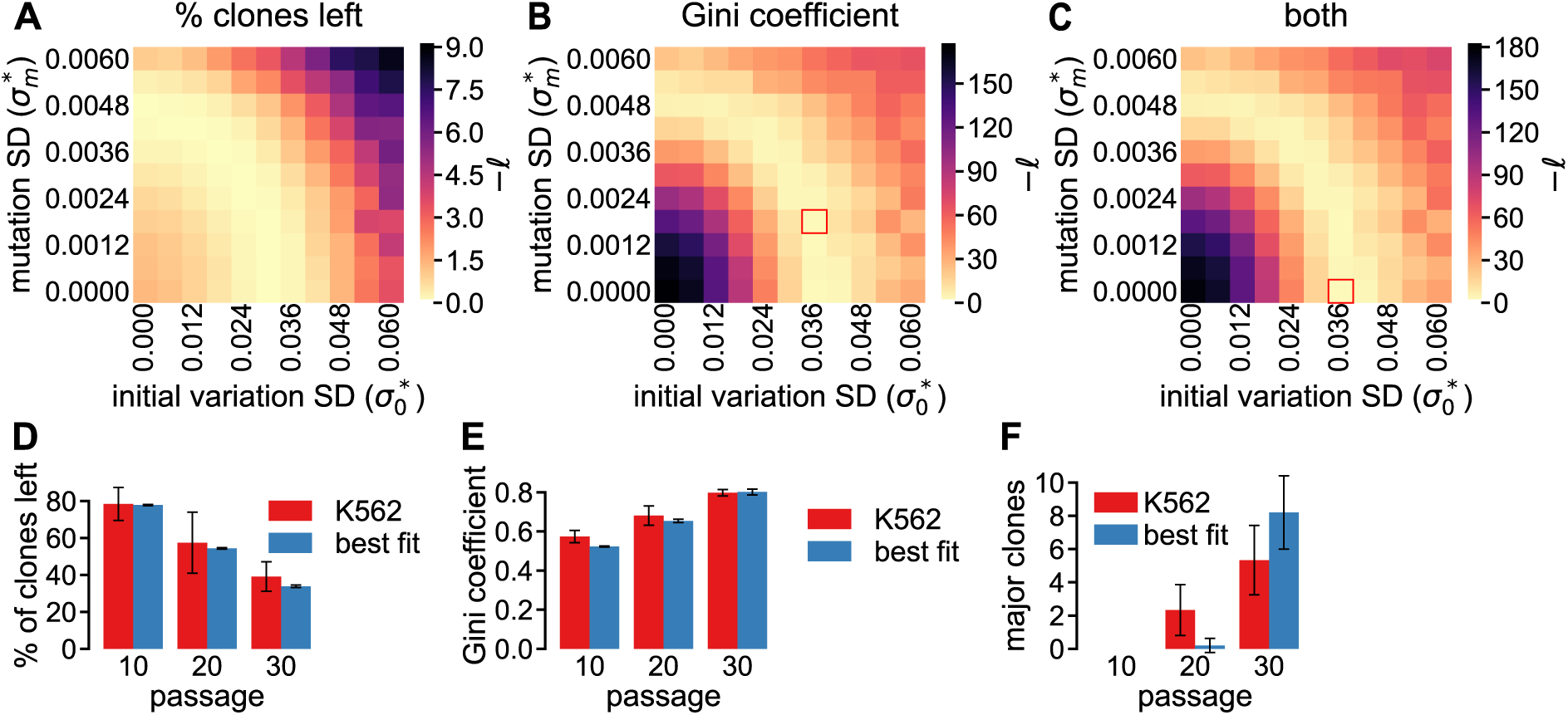
ABM simulations describing evolution of division rate variability match results for polyclonal K562 cell line. A-B Maximum Likelihood estimator ([lscript]) based on the percentage of clones lost (A), on the Gini coe cient (B), and on both metrics (C) for a range of initial division rate SDs (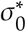) and mutation SDs (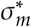). Note that we plot −[lscript] in these plots and thus its minimum value is sought. D-E comparison of clone loss (D) and clonal dominance (E) observed in simulations with the best fitting parameter values for the Gini coe cient (red rectangle in B) and in the experiments with polyclonal K562 cells. F Comparison of the number of major clones, i.e. clones representing more than 1% of the population, developing in simulations with the parameter set highlighted by the red rectangle in B and in the experiments with polyclonal K562 cells. All simulation results are the mean of 10 simulations and the results for the polyclonal K562 cells are the mean of 3 replicates, with the error bars representing the SD.

### Fitting the ABM describing division rate evolution to the K562 data

To fine-tune the model for the polyclonal K562 cell line measurements, we perform a maximum likelihood estimation (MLE) for the simulations shown in Fig 4 to identify the best fitting values for 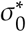 and 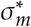. For this we consider the errors of all metrics included in the MLE to be normally distributed, so we can use the following definition of the loglikelihood:

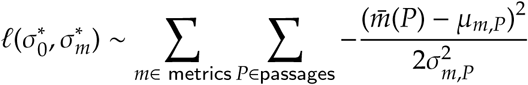

with *m* (*P*) the mean of 10 simulations for metric *m* at passage *P*, and μ_*m*_,*P* and ˙_*m*_,*P* respectively the mean and SD of the three polyclonal K562 replicates for metric *m* at passage *P*. Using passages 10, 20, and 30, and either the fraction of clones left, the Gini coe cient, or both metrics, we obtained heatmaps of the likelihood (Fig 5A-C). The simulated clone loss is quite similar to that for the polyclonal K562 cells (Fig 5A), independent of ˙^*^_*m*_ and ˙^*^_0_. This fits with the previous observation that a model with stochastic growth, but with identical division rates for all cells, already closely fits clone loss (Fig 4A). In contrast, the Gini coe cient only fits well for a defined region of parameter values (Fig 5B) and as a result this metric dominates the MLE based on both parameters (Fig 5C).

The area of best matching parameter sets also includes sets without mutation or initial division rate variation, suggesting that the source of the division rate variability cannot be determined. However, it seems most likely that both mechanisms are active because mutations before the experiment started would already induce initial division rate variation and these mutations continue to happen upon cell division. Consistent with this, the best fit corresponds to a model that includes both mechanisms (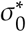 = 0.036 and 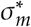 = 0.0018) and the simulation results are close to the experimentally observed Gini coe cient (Fig 5D) and clone loss (Fig 5E). Furthermore, major clones, i.e., clones representing more than 1% of the population, do develop in the ABM simulations (Fig 5E).

To test the e ect of initially present heterogeneity on clone loss and the development of clonal dominance, Porter *et al.* [13] created a monoclonal K562 cell line and showed that clone loss was reduced and clonal dominance only slightly developed (Fig. 1 F). Because this cell line was derived from a single cell, it is expected to have a lower initial variation yet similar mutation dynamics as other K562 cells. We tested this hypothesis by running a parameter sweep for 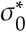 and 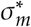 in which each simulation starts with the initial distribution of the monoclonal K562 cells.

Performing an MLE for either clone loss (Fig 6A) or the Gini coe cient (Fig 6B), we found that our model fails to produce a good fit for the clone loss (the maximum [lscript] in Fig 6A is -780), while the fit for the Gini coe cient is much better (the maximum [lscript] in Fig 6B is -40). In line with our expectations, the optimal fit of the Gini coe cient occurs at a 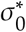 that is clearly lower than the optimal fit for the polyclonal K562 data (cf. Fig 5B). Moreover, the best-fitting 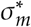 values for mono- and polyclonal K562 data are similar.

Although the optimal fitting parameters for the Gini coe cient show a close fit for the Gini coe cient (Fig 6C), this same parameter set produces a clone loss that follows a similar pattern as the monoclonal K562 cells, but the simulated clone loss is much higher (Fig 6D). However, the overestimation of clone loss by our model may actually be an underestimation of *in vitro* clone loss that is quantified by the PCR and sequencing procedure. Because any clone that had a barcode in the reference library is counted, so-called *spurious reads* may cause false positives and thus underestimate actual clone loss. These reads are at least partly the result of systematic errors that are common with next-generation sequencing methods [29]. Porter *et al.* [13] elegantly showed that the sequencing results were reproducible by pair-wise comparing the clone frequencies in four samples of the plasmid library. Nevertheless, this analysis does not completely rule out spurious reads because read errors are correlated to specific base sequences [30] and such errors are expected to occur at similar frequencies in each replicate [31]. To test the e ect of spurious reads we artificially ‘contaminated’ the clone sizes returned by the simulations with spurious reads (see details in S1 File). This analysis showed that such spurious reads can decrease clone loss to such an extent that the model matches *in vitro* clone loss. The e ect of spurious reads on the Gini coe cient was less pronounced, arguing in favour of fitting to the Gini coe cient rather than to clone loss data or to a combination of the two metrics. Furthermore, results were similar for the HeLa cells, i.e. for that cell line clone loss was not matched well whereas the Gini coe cient and the number of major clones was (for details see S2 File). Altogether, these results indicate that variation in division rates explains the dynamics of the clone size distribution described by Porter *et al.* [13].

**Figure 6:**
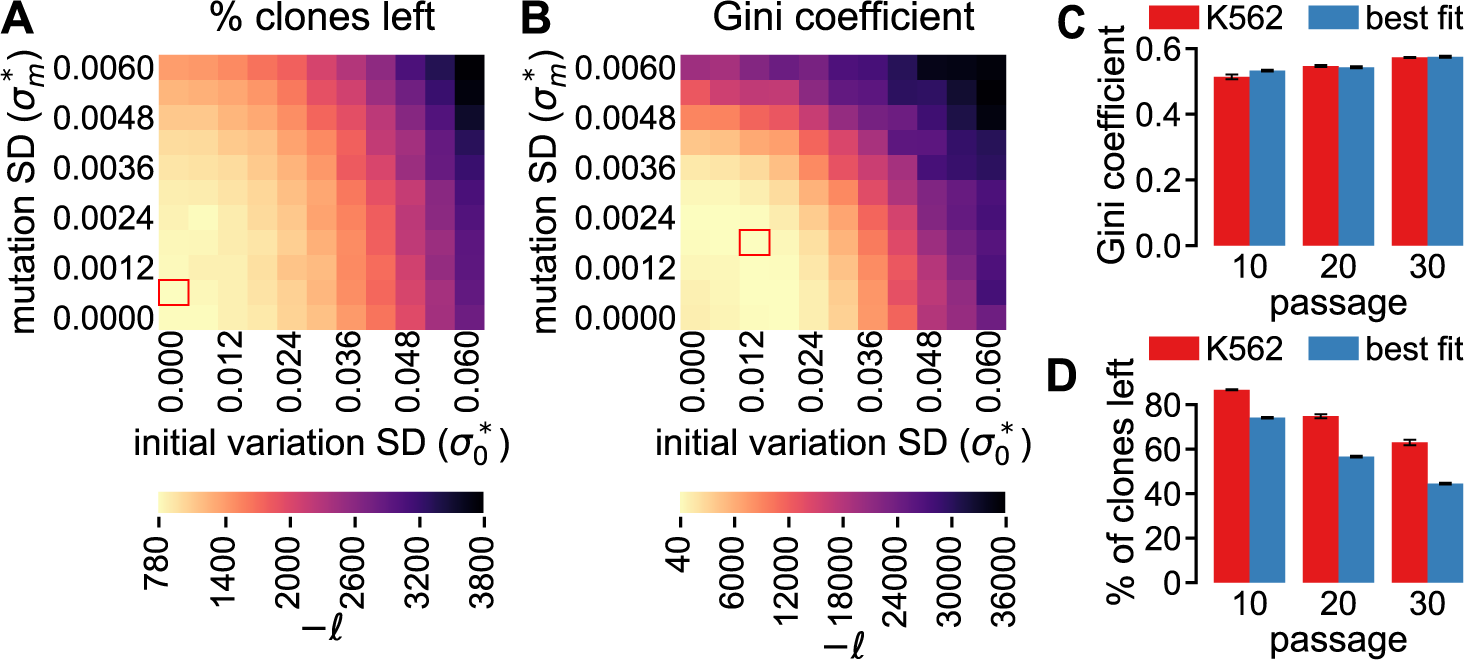
ABM simulations match the limited clonal dominance development for the monoclonal K562 cell line. A-B Maximum Likelihood estimator ([lscript]) based on clone loss (A) or Gini coe cient (B), for a range of initial division rate SDs (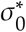) and mutation SDs (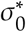). C-D Clone loss (C) and clonal dominance (D) in simulations, with the parameters from the red rectangle in B, and in the experiments with monoclonal K562 cells. All simulation results are the mean of 10 runs and the results for the monoclonal K562 cells are the mean of 3 replicates, with the error bars representing the SD.

## Discussion

Employing a simple model of stochastic cell division and passage, we showed that most of the clone loss taking place during passage, observed by Porter *et al.* [13], is due to the random loss of clones during passage. In contrast, the progressive clonal dominance that developed in the same experiments, cannot at all be explained by random clone loss. Extending the simulations with indefinitely dividing CSCs and DCs with limited division capacity did not induce progressive clonal dominance and led to a much larger clone loss than observed experimentally. However, a model in which variability of division rates was present and was subject to evolution resulted in a progressive clonal dominance matching the dominance observed experimentally. The level of correspondence between clone loss and the simulations varied per tested cell line, which we hypothesize to result from errors during sequencing as these are known to occur frequently and at a reproducible rate [29,31]. We showed that indeed such errors have a much larger e ect on the measured clone loss than on the Gini coe cient, which we used to characterize clonal dominance. Altogether, our model provides strong evidence that heritable division rate variation rather than the presence of CSCs induces the changes in clonal distribution observed by Porter *et al.* [13]. Note that our model was set up to mimic *in vitro* growth and cannot be directly extrapolated to predict the development of heterogeneity within real tumors, because in that case spatial e ects such as contact inhibition of growth and signals from the tumor micro-environment are likely to play a role. Nevertheless, our simulations elucidate which e ect evolution of division rates can have on the clonality of a cell population, and such e ects are likely to also be important *in vivo*. For example, several *in vivo*, microscopy-based, lineage tracing studies [10,11], ascribe the development of large monochromatic patches from a mosaic pattern to CSCs. Based on these observations, Driessens *et al.* [10] proposed a mathematical model with CSCs and DCs that closely fitted the clone sizes they observed in their experiments with skin papillomas. However, their model considered CSCs to divide twice as fast as DCs, while CSCs are typically thought to divide slower or at a similar rate as DCs [4]. It would therefore be interesting to investigate whether division rate variability could also contribute to such *in vivo* lineage tracing data.

Although there is in general ample evidence for the existence of CSCs [1,3,4], our model results do not point to a role for CSCs in the experiments of Porter *et al*. Indeed, in the CSC growth model with division rate heterogeneity, clonal dominance appeared in combination with a massive clone loss. This clone loss occurred because only clones that had a CSC at initialization had any chance of generating o spring in the long term, and even those clones could disappear when CSCs were accidentally lost during passage. Consistent with this explanation, removing the distinction between CSCs and DCs in our model, led to clone loss closely matching the experimental observations. Recent studies showed that the CSC fate is plastic, meaning that differentiated cancer cells sometimes can become CSCs [4,32]. This CSC plasticity might provide an alternative mechanism to prevent clone loss, by enabling clones to (re)acquire CSCs. Adding such plasticity to our CSC growth model would give all clones the potential to generate offspring indefinitely, making it similar to a model in which all cells divide indefinitely, and should thus be able to fit *in vitro* clone loss. To test if such a model with clonal evolution of division rates would develop clonal dominance we extended the CSC model with heritable heterogeneous division rates (see Methods). Simulation with this model indeed showed that clonal dominance develops (see Fig S3 Fig). However, in order to limit clone loss, it was required to have a high probability of symmetric CSC division and a large fraction of initial CSCs in the simulations. Therefore, we expect that a model with CSC plasticity can only reproduce the *in vitro* findings when the transition from DC to CSC is common.

Reproducing the *in vitro* results in our simulation was possible when we considered the division rates of tumor cells to vary between clones and to be fully and directly inherited from the parent cell (although the rate could be changed by mutation). Whereas classical studies have provided ample evidence for division rate heterogeneity among tumor cells [33–35], we are unaware of observations of its full and direct inheritance. Nevertheless, Gray *et al.* [33] showed that when melanoma cells are iteratively grown in mice, isolated, and transferred into new mice, the tumor growth speed increased every generation, which indicates that fast dividing cells generate offspring that also divide fast. More recent work further supports the assumption of heritable division rates by showing a strong, positive, correlation in the division rate of B-cell siblings [36]. However, other studies have shown that the division times of breast cancer cells correspond less between parents and offspring than between siblings [37], and that in lymphoblasts the division times of parents and offspring do not correlate at all [38]. These findings indicate that the child’s division rate is not a direct copy of the parent’s division rate. Sandler *et al.* [37] propose a *kicked cell cycle* model where the cell cycle length is determined by the level of an oscillating protein which is inherited from the parent and the phase of this protein determines the time between birth and division. Hence, such a model results in similar division times for siblings, while the correlation between division times of parents and offspring depends on the cell cycle duration [37]. These observations show the need for a better understanding of how the division rate of child cells depend on the parent, which can be achieved from lineage tracing studies employing imaging of multiple divisions over time.

While our model explains the strong clonal dominance evolving over time *in vitro*, it does not perfectly match all *in vitro* observations of Porter *et al.* [13]. The main discrepancy is that our model overestimates the clone loss observed with monoclonal K562 cells or HeLa cells (and is also slightly off for polyclonal K562 cells). Clone loss is minimal when we omit all division rate variation (Fig 2A), but even in such simulations clone loss is overestimated. Clone loss can also be reduced by increasing the percentage of clones that is passed (S4 FigA), however this also reduces clonal dominance (S4 FigB). Alternatively, it could be that passaging occurs with a bias for small clones to survive passaging, for instance when clones closely attach to each other *in vitro* and small groups of attached cells are more likely to be selected for passaging than large groups. However, this explanation seems unlikely considering that trypsinization was su cient to detach any adhering cells. Therefore, we consider it likely that *in vitro* clone loss was actually underestimated in the experimental data due to the presence of spurious reads. Our analysis (S1 File) confirmed that our model would better fit the *in vitro* data if such spurious reads are included in the simulation results.

Altogether, in this work we used a computational approach to test di erent hypotheses for the development of clonal dominance and showed that only the presence and evolution of division rate heterogeneity, can reproduce the experimental observations. Hence, this study showcases the value of computational modeling in the interpretation of experimental results. In the future, the model could be further extended to improve its power, especially for comparison with *in vivo* data. For this, the model should be extended with an explicit representation of space and physical interactions between cells [39]. With such a model it becomes possible to explore the consequences of division rate variability while comparing with intra-vital images studies.

## Methods

### Analysis of sequencing data

The sequencing data were downloaded from the NIH Sequence Read Archive (https://www.ncbi.nlm.nih.gov/sra/SRX535233) using the SRA Toolkit. The downloaded FASTQ files were processed with our own code (S3 File), which collects all barcodes for which the read quality is 56 or higher and counts the occurrence of each barcode. We used our own code rather than the ClusterSeq code (i.e., the code developed and used in Porter *et al.* [13]), because ClusterSeq clusters barcodes for each dataset separately, which may result in identical barcodes ending up in a di erent cluster at di erent time steps of the same experiment.

To generate a reference library for the cell lines barcoded with the lentiviral vector, we merged the four plasmid library samples, excluding any barcode with a frequency smaller than 0.0002% or that appeared in only one biological replicate. The resulting reference library, containing 13325 barcodes, was then used to select only these known barcodes from the experimental data (S1 Dataset). Finally, we processed the FASTQ files containing the reads for the experiments with the polyclonal K562, monoclonal K562, and HeLa cell lines. The full processing of these files is outlined in S3 File.

### SSA model of stochastic growth & passage

In order to understand the clone size dynamics of tumour cells, we employed Gillespie’s Stochastic Simulation Algorithm (SSA) [26] to simulate the growth and passage of a cell population. The SSA allows to follow the size *N i* of each clone *i* over time rather than that of single cells and we applied this to the polyclonal K562 cell line. The simulations were initialized based on the read counts for this cell line at passage 0 (see S2 Dataset), which contains the counts for *n*library = 13325 unique barcodes. Hence, we initialized *N_i_* for 1 *i n* library as 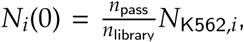 where *N*_K562,*i*_ is the read count for clone *i* in the polyclonal K562 sample and *n* pass = 3 • 10^5^ is the number of cells that are selected during passaging (i.e., also for the initial passage).

The simulations closely follow the experimental procedure of iterated growth and passage: the initial population of 3 • 10^5^ cells grows until the critical population size *n* crit = 4 • 10^6^ is reached, then *n* pass cells are selected randomly and taken to the next generation. We implemented the SSA using the τ-leaping algorithm [40], in which size *N_i_* of clone *i* is described by:

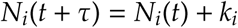

with time step τ, and *k_i_* taken from a Poisson distribution with mean *rN i* (*t*)τ, where *r* denotes the division rate. These τ leaps are performed until the population size 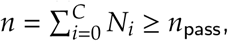, where *C* is the number of clones. We set τ = 0.0005, which resulted in a simulation time of typically Ȉ100 seconds (for Ȉ12,000 clones with one species and *r*=1) with similar results as for lower τ values (S5 Fig). The source code of the simulations can be found in S4 File.

To test how CSC driven growth a ects the clone size evolution, we incorporated a previously published model of CSC driven growth [28]. In this model CSCs proliferate at a division rate of *r*CSC and division can result either in two CSCs with probability *p_1_*, in a CSC and a DC with probability *p_2_*, or in DCs with probability *p_3_* (Fig 3A). DCs proliferate at a division rate *r*DC until they reach their maximum number of divisions *M* and then, following Weekes *et al.* [28], they die with a rate *r*DC. For simplicity, we did not consider random cell death of CSCs and DCs, because this process only a ects the population division rate. This division scheme is incorporated in the growth model by defining for each clone *i* the number of CSCs *N* _CSC,*i*_ and the number of DCs *N* _DC*m*,*i*_ of age *m* (where *m* can take values ranging from 0 to the maximum age *M*). Then, for each clone *i* there are 5 possible transitions:

1. CSC → 2CSC: *N_CSC,i_* (*t* + τ) = *N_CSC,i_* (*t*) + *k_i_*;
2. CSC → CSC + DC: *N_DC_0_,i_* (*t* + τ) = *N* DC_0_,*i* (*t*) + *k_i_*;
3. CSC → 2DC: *N_CSC,_* (*t* + τ) = *N_CSC,i_* (*t*) − *k_i_* and *N_DC_0_,i_* (*t* + τ) = *N_DC_0_,i_* (*t*) + 2*k_i_*;
4. DC_*m*_ → 2DC_*m*_+1: *N_DC_m_,i_* (*t* + τ) = *N_DC_m+1_,i_* (*t*) − *k_i_* and *N_DC_m__* 0 ≤ *m* < *M*;
5. DC_*M*_ →: *N_DC_M_, i_* (*t* + τ) = *N_DC_M,i__* (*t*) − *k_i_*,

where *k_i_* is obtained as described above, except for transitions 1-3 where the CSC division rate is multiplied by the respective transition probability. At initialization, the number of CSCs in each clone is set to *N* CSC, *i* = CSC_0_*N i*, rounded to the nearest integer. The remaining cells are distributed evenly over the *M* DC species, while rounding to the lowest integer: *N_DC,i_* = 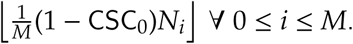. When, due to rounding, 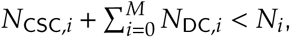, the remaining cells are randomly distributed over all species of clone *i*.

To test the e ect of division rate heterogeneity on the development of the clone size distribution, the division rates *r*i,CSC and *r*i,DC of each clone *i* are multiplied by a randomly chosen value *X_i_* that is obtained from a normal distribution with mean 1 and SD 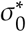, and *X_i_* is set to 0 when *X_i_* <0. Note that 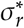 is the division rate SD relative to the mean division rate; the actual division rate SD is 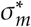 • *r*. Because *X_i_* is linked to clone *i*, the division rates are inherited upon division.

### ABM of stochastic growth & passage

The agent based model (ABM) differs from the SSA model in the explicit representation of individual cells rather than of clones. In the ABM, each cell *i* has a barcode *b_i_* and a division rate *r_i_*. All simulations are initialized based on sequencing data of either the polyclonal K562 cells, the monoclonal K562 cells, or the (polyclonal) HeLa cells, which can all be found in S2 Dataset. First, we create a *master population* from which the initial population of each replicate is selected. The master population consists of 5 • 10^6^ cells, each of which contains a barcode following read abundance in the sequencing data, such that the relative abundance of the barcodes is initially correct. Then, a division rate *r_i_* = *r*_0_ max(*X*, 0) is assigned to each cell in the *master population*, where *X* is randomly drawn from a normal distribution with mean 1 and SD 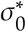, and *r*_0_ is the initial mean division rate. The seed of the random number generator used for division rate selection is kept constant across simulation replicates such that the same division rate is associated with the same cell and the same barcode in every replicate. Finally, *n*_init_ cells are taken from the initial population to be used in the simulation of iterated growth and passage. Note that the initialization omits the clonal dynamics during the 8 to 9 days before the growth & passage experiments started. During this time interval, a within-clone correlation of the division rates likely develops, which is missing in the simulated cell population. As a result the model does not reproduce the overlap between major clones that was observed by Porter *et al.* [13].

Division is modeled using the dynamic Monte Carlo method [41] implemented with a first reaction method. Whenever a cell is created, either during initialization or after division, the next division time for that cell is assigned according to 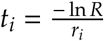, where *r_i_* is the cell’s division rate and R is a random number from a uniform distribution between 0 and 1. Then, by creating an ordered list of next division times, population growth can be simulated e ciently. When division occurs, a new division rate *r_c_* is assigned to each newly created cell according to *r_c_* = *r_p_* max(*Y*, 0) with *r_p_* the parent’s division rate and *Y* taken from a normal distribution with mean 1 and SD 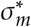. Note that 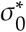 is the initial division rate SD relative to *r*_0_; the actual SD of the initial division rates is 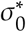 • *r*_0_. As in the SSA, division continues until the population size reaches its maximum *n*_crit_ after which *n*_pass_ cells are randomly selected and passed to the next generation. The source code of the simulations can be found in S5 File.

Note that there are some implementation di erences between the aforementioned SSA model and the ABM, which result in small quantitative di erences in the simulation results. These di erences occur for two reasons: First, growth is slightly underestimated in the SSA model, and the exact underestimation depends on the clone size. Second, a clone in the ABM can include cells with di erent division rates, which is not the case in Gillespie simulations.

## Supporting information

**S1 Fig.**
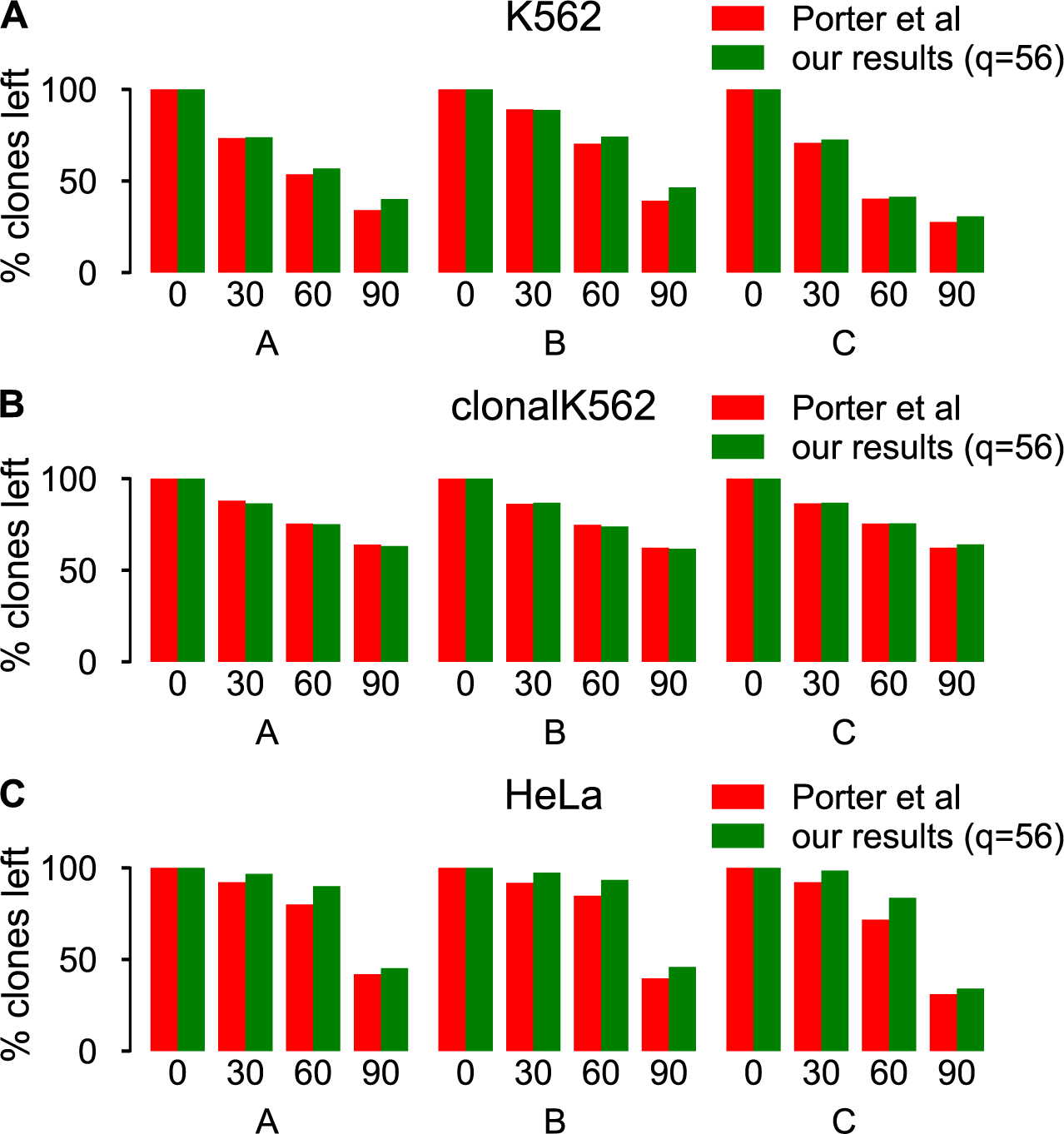
Comparison of clone number in [13] and in our analysis. A Polyclonal K562 cells. B Monoclonal K562 cells. C HeLa cells. The values in the original publication were retrieved from [13] (Figure 3f, Additional File 5d, and Additional File 12d), whereas a description of our analysis can be found in the Methods.

**S2 Fig.**
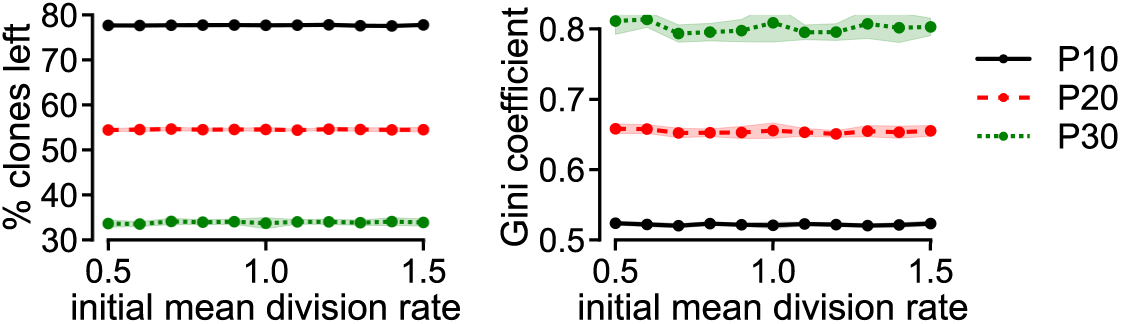
The initial mean division rate in the ABM does not a ect the clone size distribution. Clone loss (left panel) and Gini coe cient (right panel) for a series of simulations with 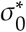 = 0.036 and 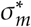 = 0.0018 and varying values of the initial mean division rate *r* 0. All data points are the mean of 10 simulations and shaded areas (only visible for the non-solid lines) represent the SD.

**S3 Fig.**
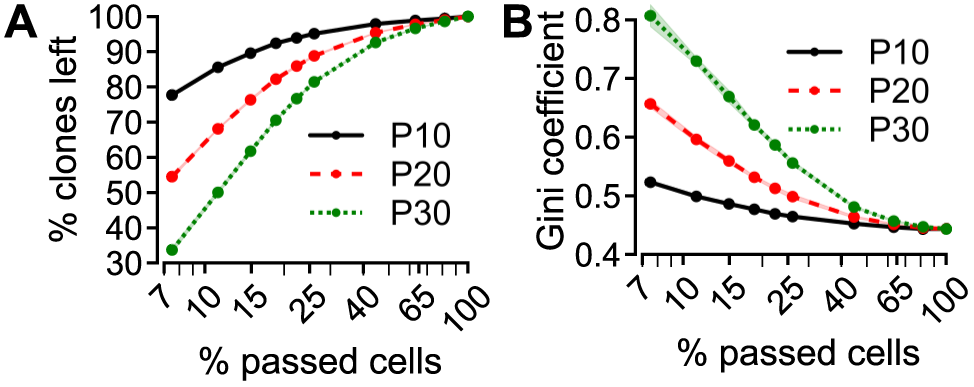
Clonal dominance develops in simulations with the CSC growth model and division rate variability. A-B Gini coe cient (A) and clone loss (B) in a simulation with 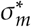 = 0.05. C-D Relation between the division rate SD and Gini coe cient (C) or clone loss (D), with the horizontal lines in C denoting the corresponding experimental values for the polyclonal K562 cell line. E Clone loss for simulations with a varying initial CSC percentage (CSC_0_) and symmetric CSC division probability (*p_1_*). The maximum of each colormap is set to the average clone loss, at the respective passage, of the three biological replicates. All values are the mean of 10 simulation replicates and the error bars (A-B) and the shaded areas (C-D) represent the SD.

**S4 Fig.**
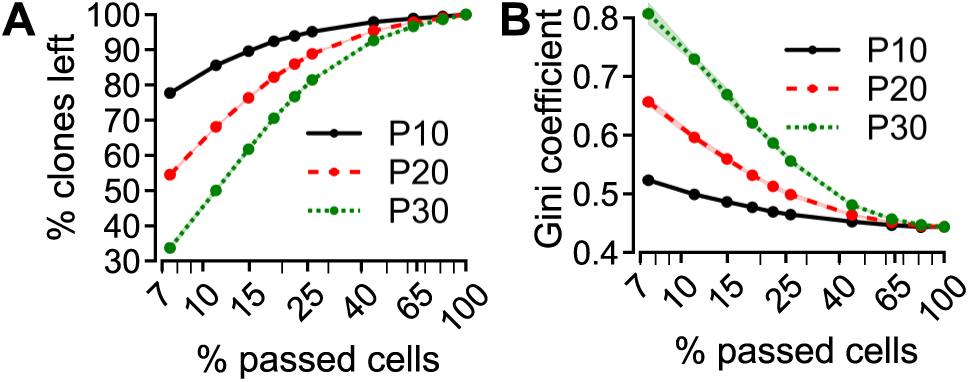
Increasing the number of cells passed reduces clone loss and clonal dominance A-B. Clone loss (A) and Gini coe cient (B) for a varying percentage of passed cells (relative to *n*_crit_). All data points are the mean of 10 simulations and shaded areas (only visible for the non-solid lines) represent the SD.

**S5 Fig.**
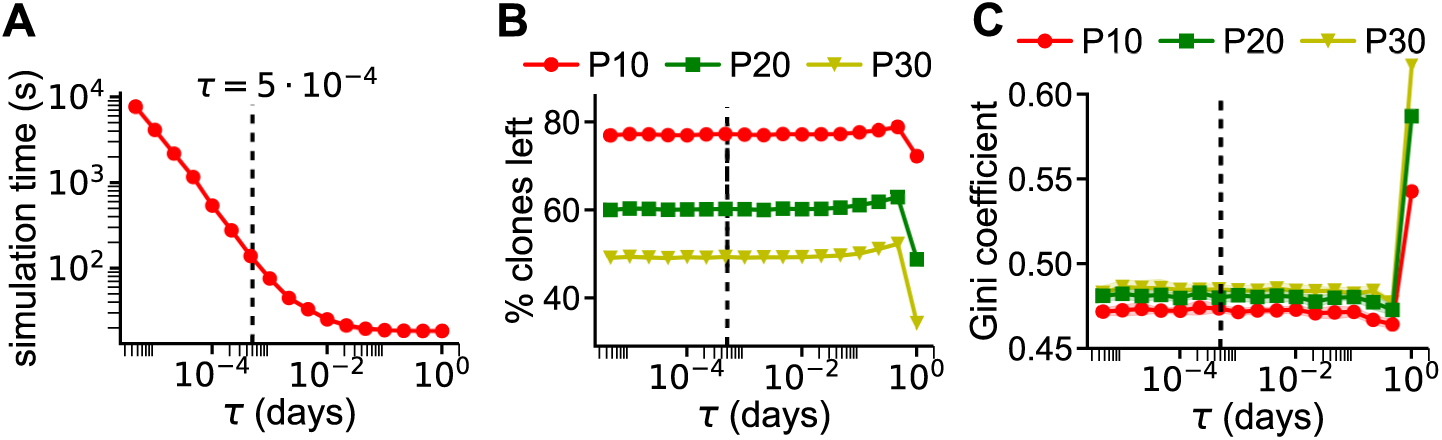
E ect of the simulation time step on the Gillespie simulations. A-C E ect of the time step (τ) on the simulation time (A), on the clone loss (B) and on the Gini coe cient(C). The vertical dashed line marks the τ used for all simulations. All points are the mean of 10 simulation replicates and the shaded areas (only visible in C) represent the SD.

**S1 File. Analysis of the e ects of spurious clones on simulated clone loss and gini coe cient.**

**S2 File. Comparison of the ABM describing division rate evolution to the HeLa data**

**S3 File. Analysis of the FASTQ files**. File contains an executable jupyter notebook and a pdf print of that notebook as well as all code needed to process the FASTQ files. *This file will be made available upon publication.*

**S4 File. Archive containing the source code for the SSA model**. *This file will be made available upon publication.*

**S5 File. Archive containing the source code for the ABM.** *This file will be made available upon publication. This file will be made available upon publication.*

**S1 Dataset. Reference library used for the analysis of the experimental data.** *This file will be made available upon publication.*

**S2 Dataset. Barcode counts of the polyclonal K562 cell line barcoded with the lentiviral vector, at passage 0**. *This file will be made available upon publication.*

## Acknowledgments

We thank Huan Yang for helpful discussions on setting up the maximum likelihood estimation.

